# Characterising stationary and dynamic effective connectivity changes in the motor network during and after tDCS

**DOI:** 10.1101/2022.09.27.509681

**Authors:** Sara Calzolari, Roya Jalali, Davinia Fernández-Espejo

## Abstract

The exact mechanisms behind the effects of transcranial direct current stimulation (tDCS) at a network level are still poorly understood, with most studies to date focusing on local (cortical) effects and changes in motor-evoked potentials or BOLD signal. Here, we explored stationary and dynamic effective connectivity across the motor network at rest in two experiments where we applied tDCS over the primary motor cortex (M1-tDCS) or the cerebellum (cb-tDCS) respectively. Two cohorts of healthy volunteers (n = 21 and n = 22) received anodal, cathodal, and sham tDCS sessions (counterbalanced) during 20 minutes of resting-state functional magnetic resonance imaging (fMRI). We used spectral Dynamic Causal Modelling (DCM) and hierarchical Parametrical Empirical Bayes (PEB) to analyse data after (compared to a pre-tDCS baseline) and during stimulation. We also implemented a novel dynamic (sliding windows) DCM/PEB approach to model the nature of network reorganisation across time. In both experiments we found widespread effects of tDCS that extended beyond the targeted area and modulated effective connectivity between cortex, thalamus, and cerebellum. These changes were characterised by unique nonlinear temporal fingerprints across connections and polarities. Our results challenged the classic notion of anodal and cathodal tDCS as excitatory and inhibitory respectively, as well as the idea of a cumulative effect of tDCS over time. Instead, they described a rich set of changes with specific spatial and temporal patterns. Our work provides a starting point for advancing our understanding of network-level tDCS effects and optimise its cognitive and clinical applications.

## 1. Introduction

Transcranial Direct Current Stimulation (tDCS) is a non-invasive neurostimulation technique widely employed to modulate cognitive and motor functions in healthy and clinical populations (Brunoni et al., 2012; Fregni & Pascual-leone, 2007). The modulation of motor processes often involves stimulating the primary motor cortex (M1) or the cerebellum (Santos Ferreira et al., 2019). It is well accepted that stimulating M1 (M1-tDCS) leads to polarity-specific changes in this area, namely excitation after anodal and inhibition after cathodal tDCS, mostly shown by motor-evoked potentials (MEP) and BOLD signal changes (Kwon et al., 2008; Nitsche & Paulus, 2000). The effects of cerebellar tDCS (cb-tDCS) are less well-understood, although there is evidence that cb-tDCS can modulate cerebellar brain inhibition (CBI), i.e., the inhibitory action the cerebellum naturally exerts over M1 (Galea et al., 2009). In particular, anodal cb-tDCS would increase inhibition towards M1 and cathodal cb-tDCS reduce it, via the dentate-thalamocortical axis (Aloi et al., 2022; Batsikadze et al., 2019; Bostan et al., 2013). This adds to the increasing evidence that tDCS effects can propagate beyond the targeted region and modulate distal functionally connected regions (Venkatakrishnan & Sandrini, 2012). For example, a number of studies on M1-tDCS indeed showed changes in other cortical areas such as SMA and premotor cortex as well as subcortical nuclei such as thalamus and putamen (Aloi et al., 2022; Jang et al., 2009; Kwon et al., 2008; Polanía et al., 2012; Zheng et al., 2011). The thalamic motor nuclei in particular play a fundamental role in motor control, as part of cerebellar and basal ganglia loops (Middleton & Strick, 2000; Parent & Hazrati, 1995). Specifically, they are thought to integrate information from cortex, basal ganglia and deep cerebellar nuclei to allow complex and fine-grained aspects of motor behaviour (Bosch-Bouju et al., 2013). Moreover, ventrolateral thalamic projections towards M1 are central for the execution of voluntary and purposeful movements, to the extent that structural impairments of this tract are associated with complete or partial absence of overt motor actions in covertly aware patients with a diagnosis of vegetative state and related conditions (Fernández-Espejo et al., 2015; Stafford et al., 2019). Being able to modulate thalamo-cortical loops can thus have a great impact over a wide variety of motor functions and clinical conditions (Caligiore et al., 2017; Carrillo et al., 2013; Fernández-Espejo et al., 2015; Joel, 2001). However, we currently lack a precise characterisation of the effects of tDCS on cortico-subcortical brain networks (Bonaiuto & Bestmann, 2015), with most available results coming from MEPs measurements or fMRI studies focusing on cortical areas only (Antal et al., 2011; Batsikadze et al., 2019; Horvath et al., 2015). Moreover, while the majority of studies deliver tDCS for 10 or 20 minutes (Reis et al., 2009), assuming that the action of tDCS is cumulative, we do not in fact know how the neural activity is dynamically modulated across time, or whether the temporal mechanisms of action depend on the specific regions that are being stimulated.

In a recent study (Aloi et al., 2022), we used dynamic causal modelling (DCM) of fMRI to demonstrate that tDCS can modulate long-range thalamo-cortical dynamics during a simple command following task involving voluntary thumb movements in response to auditory cues. Specifically, anodal M1-tDCS induced thalamic disinhibition, while cathodal cb-tDCS increased excitation from thalamus to M1 (Aloi et al., 2022) during movement blocks. Here, we build upon this by examining tDCS changes on the intrinsic brain activity at rest, when unconstrained by a task. In addition, our previous study focused on changes immediately after tDCS as compared to a pre-tDCS baseline. Indeed, this is a common focus in the literature and there is little understanding of how the effects of tDCS on complex brain dynamics evolve over time (Bestmann et al., 2015; Bonaiuto & Bestmann, 2015). In the current study, we characterise online tDCS and their dynamic changes over 20 minutes of stimulation. Specifically, we report data from two experiments that used DCM of resting state fMRI data to investigate whether M1- and cb-tDCS (respectively) are able to modulate the resting-state effective connectivity between cortical and subcortical motor areas, and how. We first explored the specific changes in neural coupling associated with short-term effects of tDCS by comparing resting-state fMRI data before and after stimulation. Then, we investigated online changes in effective connectivity at rest *during* 20 minutes of tDCS stimulation, to observe the immediate effects of ongoing stimulation. To this end, we first compared the *stationary* effects of tDCS across the 20 minutes of stimulation. Finally, we investigated the temporal dynamics of such effects using a novel sliding windows approach (Park et al., 2018) to model the network reorganisation through time.

We hypothesised anodal and cathodal M1-tDCS to respectively yield an excitatory vs inhibitory effect over M1 that would trigger further network changes in thalamocortical coupling.

Additionally, cathodal cb-tDCS would elicit inhibition over the cerebellum, consequently inducing excitation along the motor network (as in Aloi et al., 2022) with anodal cb-tDCS inducing the opposite effect. Finally, while our online analyses were exploratory due to the lack of prior literature, we expected to observe dynamic changes in effective connectivity (across timewindows) that would capture the progression of tDCS effects over time.

## 2. Materials and methods

### 2.1 Participants

We recruited a total of 49 participants across the two experiments (mean age: 25 ± 4; 15 males, 34 females) using the University of Birmingham Research Participation Scheme and advertisements across campus. All participants were right-handed according to the Edinburgh handedness inventory (Oldfield, 1971; mean handedness score: 87.2 ± 19.9 in Experiment 1; 84.5 ± 20.3 in Experiment 2). All participants were over 18 years old, and reported no history of epilepsy, migraine, neurological or psychiatric disease. They also met the eligibility criteria to enter the MRI environment and to undergo brain stimulation (we confirmed this before each individual session). We instructed participants to come to the appointment well hydrated and rested, without consuming alcohol or coffee within 24 hours prior to testing. Participants filled in the screening forms for MRI and brain stimulation and had the chance to read the information sheet and ask questions prior to providing written informed consent. They received compensation in the form of cash or research credits. The University of Birmingham’s Science, Technology, Engineering and Mathematics Ethical Review Committee provided approval for our study.

*Experiment 1* included 26 participants. Four participants withdrew before completing all the sessions, and we discarded a further participant due to corrupted data, resulting in a total of 21 participants for the analysis (age range: 18-32; mean age: 22.1 ± 3.9; 8 males, 13 females).

*Experiment 2* included 23 participants. From these, 1 withdrew from the experiment before completing all the sessions, resulting in 22 participants for the analyses (age range: 21-37; mean age: 26.9 ± 4.1; 7 males, 15 females).

### 2.2 Experimental design and procedure

Both experiments used sham- and polarity-controlled, randomised, double-blind, crossover designs, where each participant completed three tDCS sessions (anodal, cathodal, and sham) on separate days. We schedule sessions at least 7 days apart (*Experiment 1:* mean 12± 10; *Experiment 2*: mean 13± 7) to avoid carryover effects, and in a counterbalanced order across participants.

After completing pre-screening and consent procedures (see above), we set up the tDCS electrodes on the participant’s scalp in a dedicated room, before taking them to the MRI scanner. In each session (anodal, cathodal, or sham) we acquired resting-state fMRI before, during and after 20 minutes of tDCS (Figure 1).

**Figure 1.**
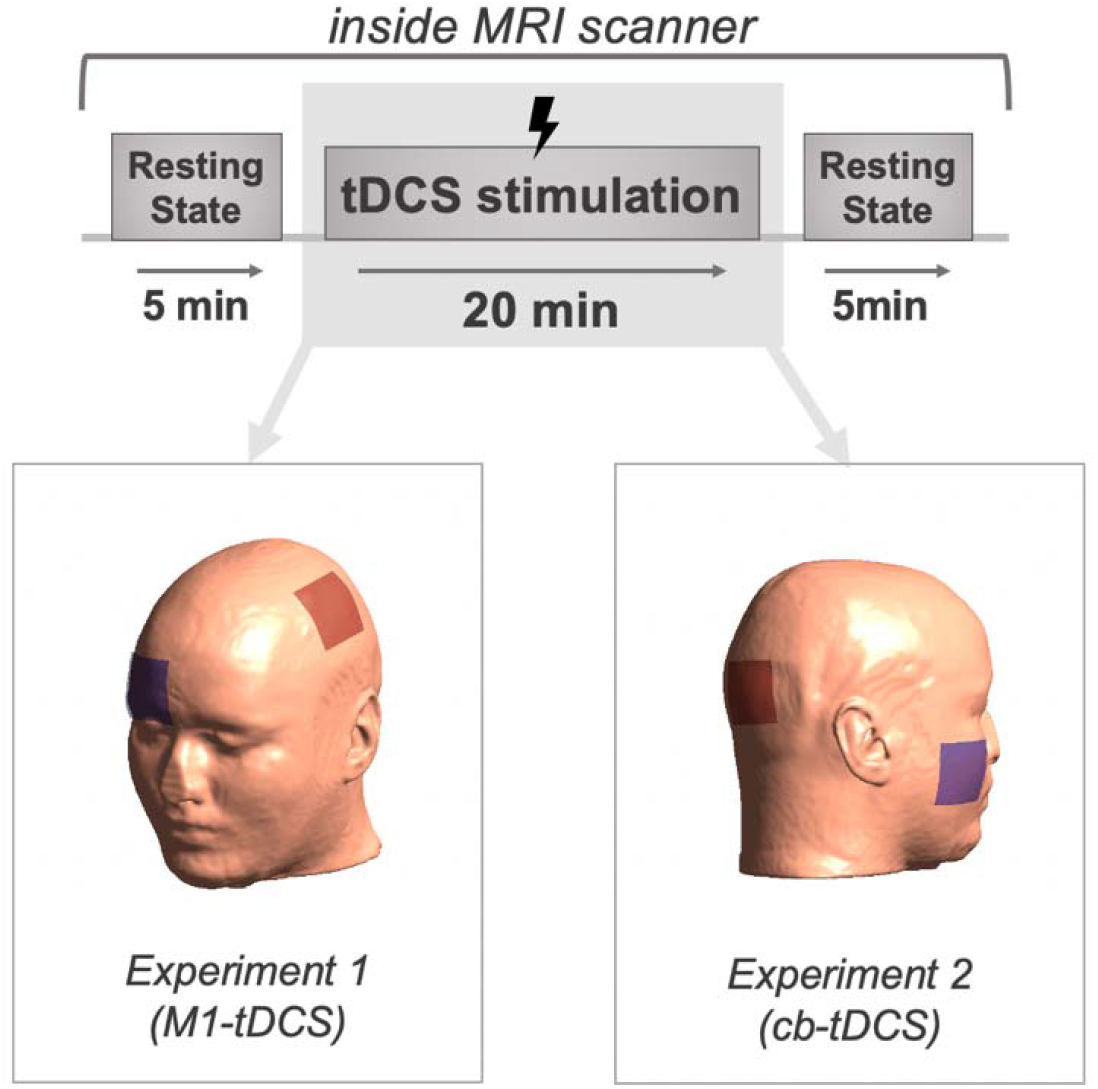
Experimental design and tDCS montages. We acquired resting-state fMRI data before (5 minutes), during (20 minutes) and after (5 minutes) tDCS stimulation (three sessions per participant – anodal, cathodal and sham stimulation – in a counterbalanced order). Bottom left: tDCS montage for *Experiment 1, with* target electrode on the left primary motor cortex (M1) and reference electrode on the right orbitofrontal cortex (OFC). Bottom right: tDCS montage for *Experiment 2*, with target electrode on right cerebellum and reference electrode on right buccinator muscle respectively.

After every session, participants completed a post tDCS perceptual scale form to report the presence and intensity of sensations and/or discomfort during stimulation. They also had to indicate whether they believed they received real stimulation or sham.

### 2.3 Electrical stimulation

We delivered the stimulation with an MRI compatible NeuroConn DC-Stimulator MR system (neuroCare Group GmbH, Munich, Germany). We used 5×5 cm^2^ electrodes with Ten20 conductive paste to increase conductivity and secured them in place using self-adhesive bandage.

#### Experiment 1

We placed the target electrode over the left M1, centred on the motor hotspot as identified by TMS prior to the first MRI session (Rossini et al., 2015), and oriented at an angle of approximately 45° from the midline. We positioned the reference electrode over the right orbitofrontal area.

#### Experiment 2

We placed the target electrode on the right cerebellar cortex (3 centimetres lateral to the inion, oriented parallel to the midline) with the reference on the right buccinator muscle (Galea et al., 2011).

See Figure 1 for a representation of both montages.

In the anodal sessions, the anodal electrode was target and the cathodal one was the reference, and we reversed this for the cathodal sessions. Half of the sham sessions replicated the anodal montage and the other half the cathodal montage. We recorded the position of the electrodes in the first session using Brainsight™ TMS neuronavigation software (Rogue Research Inc). We subsequently used the saved coordinates in session 2 and 3 to ensure consistent electrode placement.

During anodal and cathodal sessions, we delivered 20 minutes of tDCS stimulation with an intensity of 1 mA (*Experiment 1*) or 1.85 mA (*Experiment 2*) with 30 s ramp-up and ramp-down periods while in the MRI scanner. In the sham condition, we applied stimulation for 30 s only before ramping down, to emulate the sensation elicited by real tDCS (Ambrus et al., 2012; Woods et al., 2016).

### 2.4 MRI acquisition

We acquired all data on a 3T Philips Achieva scanner at the Birmingham University Imaging Centre (BUIC).

#### Experiment 1

We collected resting-state fMRI data with an EPI sequence and the following parameters: 600 volumes per run during tDCS, and 150 volumes per run for pre and post tDCS runs, 34 slices, TR = 2000 ms, TE = 35 ms, flip angle = 79.1°, matrix size = 80×80, field of view = 240×240 mm, voxel size = 3×3×3. Additionally, we obtained T1-weighted images for anatomical co-registration with parameters TR = 7.4 ms, TE = 3.5 ms, flip angle = 7°, matrix size = 256×256, field of view = 256×256, voxel size = 1×1×1.

#### Experiment 2

We collected resting-state fMRI data with an EPI sequence and the following parameters: 450 volumes per run during tDCS, 112 volumes per run for pre and post tDCS runs, 46 slices, TR = 2700ms, TE = 35ms, flip angle = 79.1°, matrix size = 80×80, field of view = 240×240 mm, voxel size = 3×3×3. T1-weighted images were also acquired with parameters TR = 7.4ms, TE =3.5ms, flip angle = 7°, matrix size = 256×256, field of view = 256×256, voxel size = 1×1×1.

In both experiments we collected other anatomical data, including diffusion weighted imaging but we did not analyse them within the current study and we will report this in separate papers. Additionally, we collected task fMRI data with a simple motor paradigm (thumb movements) before and after tDCS and we have reported this in Aloi et al., (2022).

### 2.5 fMRI preprocessing

We used SPM12 version v7771 (Ashburner et al., 2016) on MATLAB R2019b to pre-process and analyse the fMRI data. We analysed the datasets from the two studies independently but following the same pipeline, as described in here (see also Figure 2). This included standard pre-processing steps: realignment, slice timing correction, coregistration between anatomical and functional images, spatial normalisation, and smoothing with an 8mm^3^ Gaussian kernel. In addition, we applied used the TAPAS PhysIO Toolbox to model physiological noise (Kasper et al., 2017) and perform a denoising procedure that resembled the aCompCor method by Behzadi and colleagues (Behzadi et al., 2007). Specifically, we performed a principal component analysis (PCA) on *noise* regions of interest (ROIs), namely segmented masks of the white matter and cerebrospinal fluid (CSF), based on the assumption that the temporal fluctuations in those ROIs would not be related to neural activity but rather to respiratory and cardiac confounds (Behzadi et al., 2007). To create these ROIs, we used a threshold of 0.95 for voxel inclusion and eroded the masks by cropping 1 voxel. We then extracted the fMRI time series of voxels corresponding to the *noise* ROIs. The toolbox computed the first 3 principal components and returned them in the form of noise regressors. We chose the voxel inclusion threshold and the number of PCs to achieve a strict ROIs selection while keeping a sufficient number of voxels in our masks (i.e., a stricter voxel selection resulted in empty ROIs). We did not perform temporal filtering since previous studies suggested that filtering might be a redundant step if applied after denoising (Bright et al., 2017).

**Figure 2.**
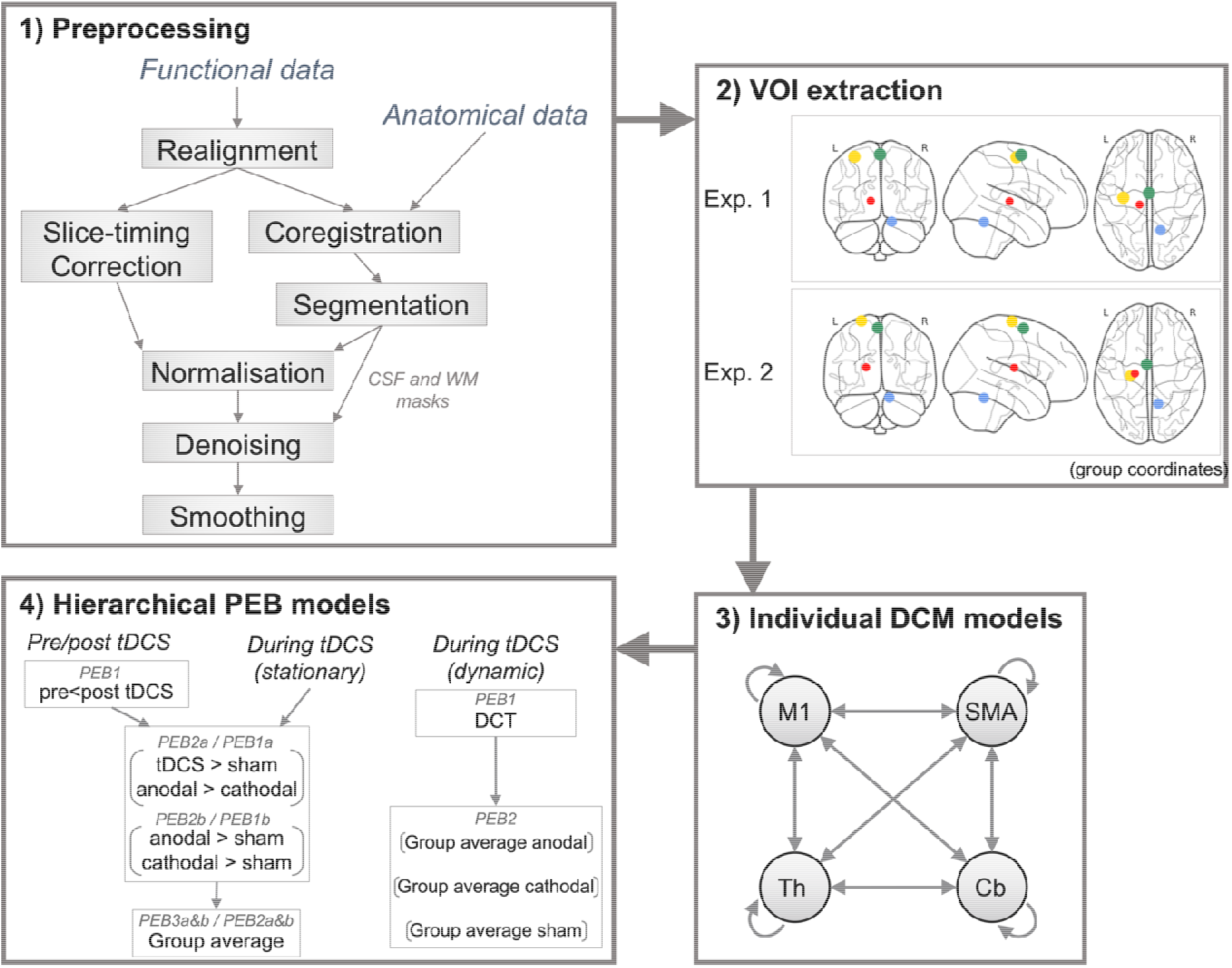
Analysis pipeline. 1) We used a standard pre-processing pipeline on SPM12, including a denoising procedure with the TAPAS Physio Toolbox; 2) We extracted time-series from 4 individually defined regions of interest: left M1, SMA and thalamus, and right cerebellum, with coordinates obtained from Aloi et al., 2022; 3) We specified and estimated fully connected individual DCM models; 4) We ran a number of hierarchical PEB models to obtain the relative group effects for the differences between pre- and post-tDCS across polarities, differences during the full run of tDCS across polarities, and dynamic temporal changes during tDCS in each polarity. These analyses were performed separately for each dataset (pre/post tDCS, full run during tDCS, sliding windows during tDCS) and *Experiment*. For stationary analyses, all group PEBs modelled the differences between tDCS and sham, between anodal and cathodal, anodal and sham, and cathodal and sham. The dynamic analysis included fitting a discrete cosine transform (DCT) with 6 basis functions across windows, then a group PEB of between-windows effects for each polarity.

### 2.6 DCM analysis

#### 2.6.1. Region selection and timeseries extraction

We extracted time-series from spherical volumes of interest (VOI) centred on functionally defined coordinates for each individual participant for the left M1, SMA and thalamus, and the right cerebellum. Specifically, we used individual peaks of activation for each region and participant for the contrast between movement and rest in a simple motor command following task where participants were instructed to move their right thumb in response to beeps, as reported in Aloi et al., 2022. We used spheres with 8-mm radius for M1 and SMA, 4 mm radius for thalamus, and 6-mm radius for the cerebellum, to account for differences in anatomical size.

#### 2.6.2. Individual level DCM specification and definition of model space

For each participant and run, we first specified individual bilinear, one-state, deterministic DCM models that fitted cross-spectral densities (as recommended for resting state data; Friston et al., 2014) to the pre-, during, and post-tDCS data. These models were fully connected across the 4 regions of interest (i.e., all self- and between region connections in the A matrix were switched on). As this is resting-state data, we set no external modulations or inputs (i.e., empty B and C matrices). At the first within-subject (and within-session) level, DCM works by coupling the specified connectivity matrix (and all possible variations of it) with a cascade of neuronal, neurovascular coupling, hemodynamics and fMRI signal models that shape the interactions between neuronal populations, cerebral blood volume, deoxyhemoglobin and observation noise in order to predict a BOLD signal from the specified neural dynamics (Zeidman et al., 2019a).

#### 2.6.3 PEB analyses

We performed three DCM analyses under the Parametric Empirical Bayes (PEB) framework for each of the two datasets independently: (1) we compared resting-state data *post-* (vs pre-) between the different tDCS polarities (and sham) to investigate short-term effects of stimulation, (2) we compared data *during* tDCS across polarities (and sham) to investigate the immediate influence of ongoing stimulation, and (3) we segmented the data *during* tDCS into sliding time-windows and fitted a Discrete Cosine Transform across them, to explore the temporal dynamics of ongoing stimulation across polarities. See Figure 2 for an overview of the analysis pipeline.

Briefly, PEB is a hierarchical Bayesian framework for group-level modelling of effective connectivity, that includes group-level constraints over the parameter estimation at the individual level, enhancing estimation precision (K. Friston et al., 2018). To find which model best explains the real observed data first *Bayesian Model Inversion* is applied to each model to calculate the parameters (connection strengths) that yield the best trade-off between maximum data explanation and minimum complexity. Then, Bayesian *Model Comparison* is used to compare the evidence (or Marginal Likelihood) for each model, first at an individual, and subsequently at a group level in a hierarchical manner. After that, *Bayesian Model Reduction (Zeidman et al., 2019b*) is used to prune away connections that do not contribute to the model evidence, by comparing evidence across reduced models each having specific parameter combinations. Finally, *Bayesian Model Averaging* (BMA) computes the average of parameters across all models weighted by the models’ posterior probability. We only report parameters that exceed 95% of posterior probability (equivalent to a Bayes factor of 3 and considered as strong evidence).

##### 2.6.3.1 Stationary effective connectivity post-tDCS

This analysis included 21 participants across 3 sessions (anodal, cathodal, sham) with 2 runs (pre-tDCS, post-tDCS) each, for a total of 126 individual DCMs for *Experiment 1*. The equivalent analysis for *Experiment 2* included 132 individual DCMs (22 participants x 3 sessions x 2 runs).

To test the effects of tDCS on the model parameters, we first defined a first-level PEB for each participant and polarity, in which we computed the difference between pre and post stimulation runs (pre-tDCS < post-tDCS), taking the DCM models as input. We then entered the results of this PEB into two separate second-level PEBs:

1. *PEB2a* investigated the polarity specific effects of anodal tDCS by computing the difference between tDCS stimulation (anodal and cathodal together) and sham (i.e., anodal tDCS + cathodal tDCS > sham), and the difference between anodal and cathodal stimulation (anodal > cathodal).
2. *PEB2b* explore the effects of each polarity individually by computing the difference between anodal and sham (anodal > sham), and between cathodal and sham (cathodal > sham).

Finally, two separate third-level PEBs (*PEB3a* and *PEB3b*) computed group effects by averaging *PEB2a* and *PEB2b* respectively across participants. Each final PEB thus modelled the following effects:

1. *PEB3a:* regressors for the main effect of time (pre<post), main effect of tDCS, interaction between tDCS and time, main effect of anodal vs cathodal, and interaction of anodal vs cathodal with time;
2. *PEB3b:* regressors for the main effect of time, main effect of anodal vs sham, interaction between anodal vs sham and time, main effect of cathodal vs sham, and interaction of cathodal vs sham with time.

As our aim was to test the effects of tDCS, we only report the outcomes of the interactions between time and polarity. See Figure 3 for a summary of the hierarchy.

**Figure 3.**
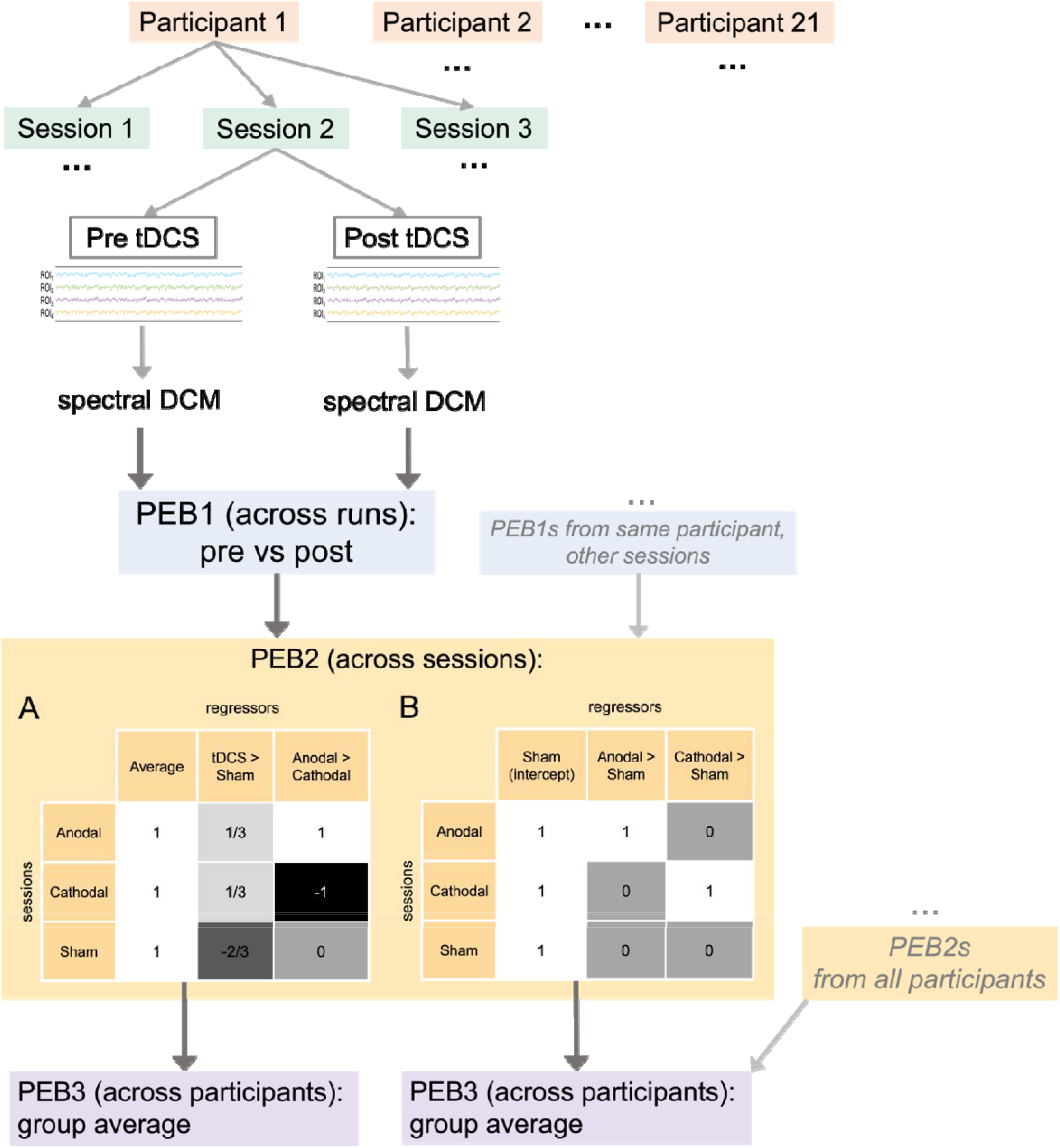
Overview of DCM and hierarchical PEB analysis of data *pre- and post-tDCS*. Sections A and B show the PEB2 regressors used to generate the contrasts of interest.

##### 2.6.3.2 Stationary effective connectivity during tDCS

This analysis included in 63 individual DCMs (21 participants x 3 sessions) in *Experiment 1*, and 66 in *Experiment 2* (22 participants x 3 sessions), each modelling effective connectivity during tDCS.

Similarly to above, we first computed two separate PEBs for each participant:

1. *PEB1a* encoded the difference between tDCS stimulation (anodal and cathodal together) and sham, and the difference between anodal and cathodal stimulation.
2. *PEB1b* computed the difference between anodal and sham (anodal > sham) and between cathodal and sham (cathodal > sham).

Two second-level PEBs averaged commonalities across participants for PEB1a and PEB1b respectively (Figure 4).

**Figure 4.**
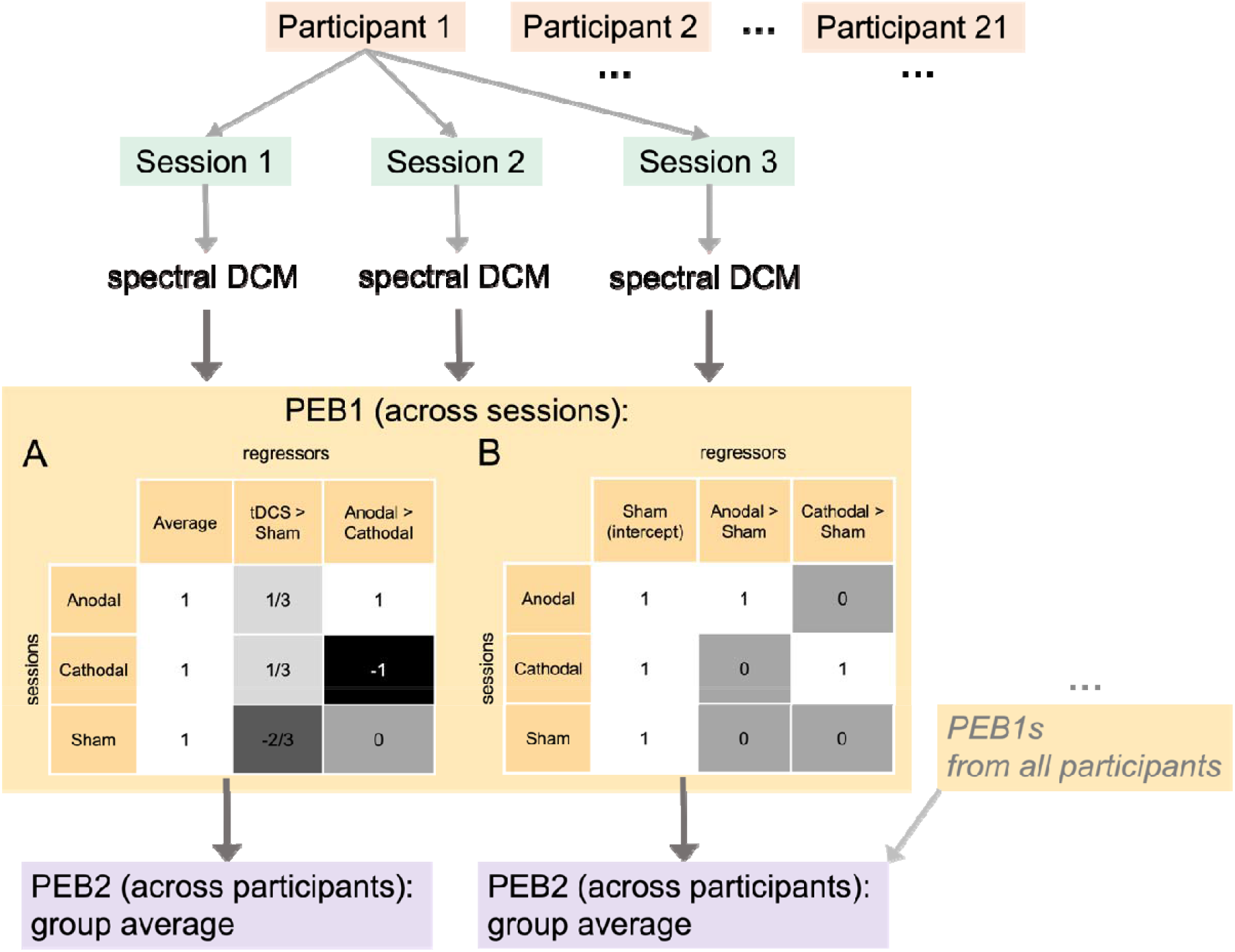
Overview of DCM and hierarchical PEB analysis of data *during* tDCS (stationary). Sections A and B show the PEB1 regressors used to generate the contrasts of interest.

##### 2.6.3.3 Dynamic effective connectivity during tDCS

Here, we aimed to study changes in DCM dynamics across time-windows in response to *online* tDCS. First, we built individual DCMs for overlapping time windows on the data acquired during stimulation. Each time-window had 200 s duration and a 50% overlap with the subsequent window (100 s), as previously done in Park et al., 2018), resulting in 11 sliding time-windows per session. This produced a total of 693 individual DCM models (11 time-windows x 3 sessions x 21 participants) in *Experiment* 1 and 726 individual DCM models (11 time-windows x 3 sessions x 22 participants) in *Experiment 2*.

A first-level PEB consisting of a set of temporal basis functions (Figure 5, section C) modelled the effective connectivity fluctuations across time windows (Park et al., 2018; Van de Steen et al., 2019). Specifically, we employed a discrete cosine transform (DCT) with 6 functions, where the first function computed the average, and the other 5 modelled different fluctuation rates across windows. This included fluctuations between 0.0078 Hz and 0.1250 Hz, within the range of canonical resting-state spontaneous fluctuations (Auer, 2008). DCT is commonly used to describe the spatial and temporal properties of resting-state functional connectivity patterns (Fransson, 2005), and has been previously used to investigate the dynamic effective connectivity of resting-state fMRI with the sliding-windows approach (Park et al., 2018).

**Figure 5.**
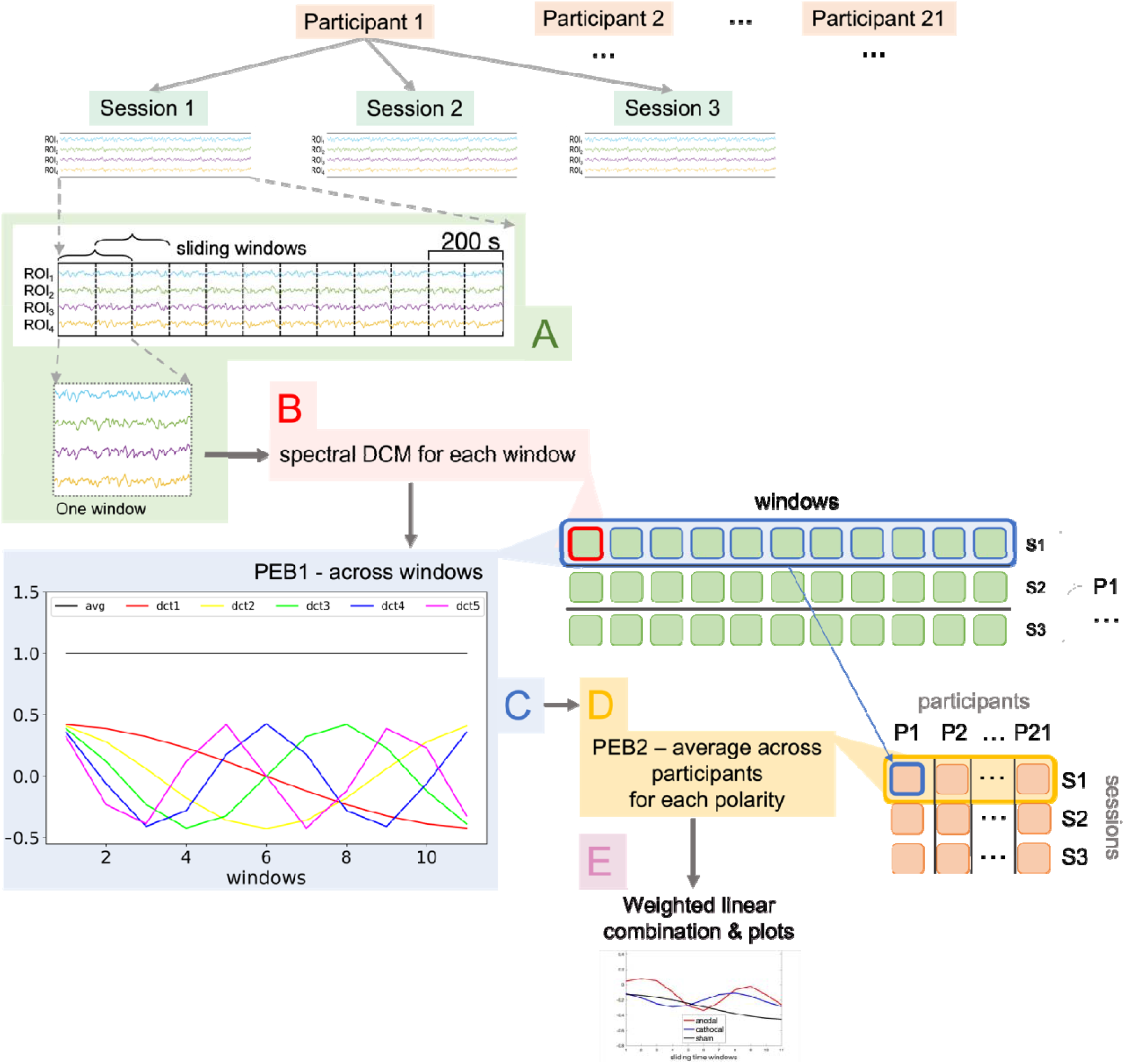
Overview of DCM and hierarchical PEB analysis of data *during* tDCS (sliding windows approach). (A) We divided each of the three sessions undertaken by each participant into 11 sliding time-windows with 200 s duration and 100 s overlap. (B) We then specified individual DCM models for each time-window. (C) In the first-level PEB (PEB1), we applied a discrete cosine transform (DCT) to each individual tDCS session to model the temporal dynamics between windows. The plot shows the DCT which consists of an average across windows and 5 cosine functions that oscillate at different frequencies (from 0.0078 Hz to 0.1250 Hz). (D) A second level PEB (PEB2) then computed the group effects for each tDCS polarity to allow informative data visualisation. (E) We obtained plots for each connection across participants after performing a linear combination of each cosine function weighted by its estimated parameter.

We then averaged the resulting PEBs across participants for each polarity separately to obtain the group effects of anodal, cathodal, and sham stimulation (PEB2). Each of these second-level PEBs returned a parameter matrix for each basis function, depicting which nodes of the network followed the rate of change described by the function (i.e., nodes with posterior probability > 95%), and to what extent (i.e., effective connectivity parameters). Additionally, to produce plots representing changes in connectivity across time, we performed a weighted linear combination of the basis functions for each connection under each tDCS polarity using the estimated parameters as weights, as follows:

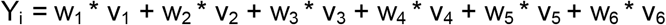

where *v* are the functions that belong to the DCT, *w* are the estimated parameters associated with them, and *i* is the polarity.

See Figure 5 for a summary of the analysis hierarchy.

### 2.7 Blinding

In this study we analysed the responses to one of the questions in the post-tDCS questionnaire (*“Did you receive REAL tDCS? If you believe that you received real tDCS, please select ‘YES’*. *If you believe that you did not receive real tDCS (i.e., received the placebo version) please select ‘NO’”*) to assess whether our blinding procedure was successful. We grouped the results in real tDCS (anodal and cathodal together) and sham stimulation, then performed a McNemar’s test to check whether participants could determine if they were receiving real or sham tDCS.

## 3. Results

### 3.1 Stationary effective connectivity pre-vs post-tDCS

#### Experiment 1 – Effects of M1-tDCS

The individual DCM models from this analysis explained an average of 87.99% of variance (SD = ±3.51; see all results from the explained variance analysis in Figure S1, Supplementary Material).

Anodal M1-tDCS reduced thalamic self-inhibition as compared to both sham and to cathodal tDCS. Similarly, it increased excitation from SMA to cerebellum and from cerebellum to thalamus also compared to both cathodal tDCS and sham. In addition, anodal tDCS also increased inhibition from thalamus to SMA and cerebellum, and excitation from SMA to M1, although these changes were not polarity specific (i.e., they were only significant in comparison to sham).

In contrast, cathodal stimulation inhibited the coupling from M1 to thalamus but increased excitation from thalamus to M1 both as compared to sham and as compared to anodal stimulation. In addition, it reduced cerebellar self-inhibition compared to sham only.

Both anodal and cathodal tDCS increased excitation from SMA to thalamus in a similar way compared to sham stimulation, whereas they induced opposing effects over the cerebellum-SMA coupling, with anodal tDCS decreasing and cathodal tDCS increasing excitation both as compared to sham and to each other. Figure 6A represents the results after M1-tDCS (see supplementary Figure S2 A for a more detailed representation including effective connectivity and posterior probability values).

**Figure 6.**
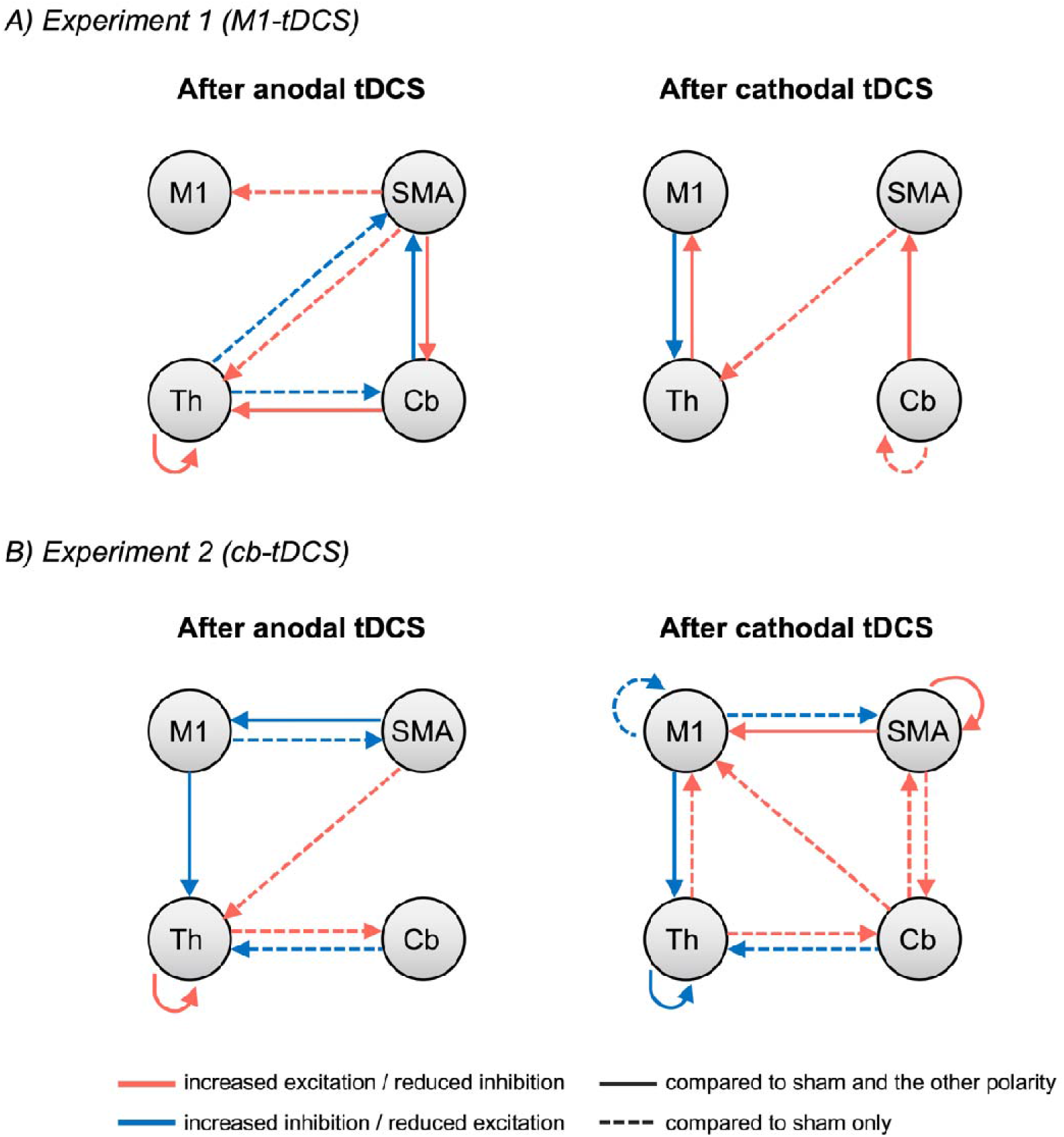
After effects of tDCS on stationary effective connectivity. The figure shows connections that exceeded the 95% posterior probability threshold in the analysis of data after tDCS compared to baseline (interactions between time and polarity only) for *Experiment 1* (A) and *Experiment 2* (B). The left and right panels represent changes after anodal and cathodal stimulation respectively. Red arrows indicate connections positively related to the regressor (i.e., more excitatory / less inhibitory as a product of the comparison), while blue arrows signal connections with a negative relation to the regressor (i.e., more inhibitory/less excitatory) as a product of the comparison). Note that self-connections are always inhibitory and thus red indicates a reduction in inhibition rather than an excitatory role per-se. Solid lines represent polarity-specific effects (i.e., effects that are significant as compared to both sham stimulation and the other polarity) while dashed lines indicate significant effects as compared to sham stimulation only.

#### Experiment 2 – Effects of Cb-tDCS

The individual DCM models from this analysis explained an average of 88.65% of variance (SD = ±4.16).

Both anodal and cathodal cb-tDCS increased inhibition from cerebellum to thalamus and excitation from thalamus to cerebellum as compared to sham (with no significant differences between polarities). In addition, they both increased inhibition from M1 to thalamus, although this was significantly stronger for cathodal stimulation. Finally, they both increased inhibition from M1 to SMA as compared to sham (with no differences between polarities).

In contrast, the two polarities had opposite effects on thalamic self-inhibition, with anodal reducing and cathodal increasing this, both as compared to sham and to each other. Similarly, they respectively decreased (anodal) and increased (cathodal) excitation from SMA to M1 (also as compared to sham and to each other).

As compared to sham only, cathodal cb-tDCS increased excitation from cerebellum to M1 and SMA, from SMA to cerebellum and from thalamus to M1, and increased self-inhibition in M1. It also decreased self-inhibition in SMA as compared to both anodal and sham stimulation. In turn, anodal tDCS increased excitation from SMA to thalamus as compared to sham only.

Figure 6B represents the results after cb-tDCS (see supplementary Figure S2 B for a more detailed representation).

### 3.2 Stationary effective connectivity during tDCS

#### Experiment 1 – Effects of M1-tDCS

The individual DCM models from this analysis explained an average of 88.25% of variance (SD = ±2.96).

When analysing the full 20 minutes of stimulation under stationarity assumption, anodal and cathodal M1-tDCS elicited an opposite effect over the thalamic self-connection, whereby anodal increased and cathodal decreased inhibition, both when compared to sham and to each other. In contrast, both polarities had similar inhibitory effects on the cerebellum-SMA coupling (although this was significantly stronger for anodal). Similarly, both polarities increased cerebellar self-inhibition as compared to sham (this time with no differences across polarities) Anodal M1-tDCS also increased excitation from thalamus to cerebellum, inhibition from cerebellum to M1, and SMA self-inhibition, compared to both cathodal and sham stimulation. In addition, it increased excitation from M1 to cerebellum and inhibition from cerebellum to thalamus but these effects were not polarity specific and only appeared as compared to sham. In turn, cathodal stimulation increased inhibition from M1 to thalamus both as compared to anodal stimulation and sham. Figure 7A represents the results during M1-tDCS (see supplementary Figure S3 A for a more detailed representation including effective connectivity and posterior probability values).

**Figure 7.**
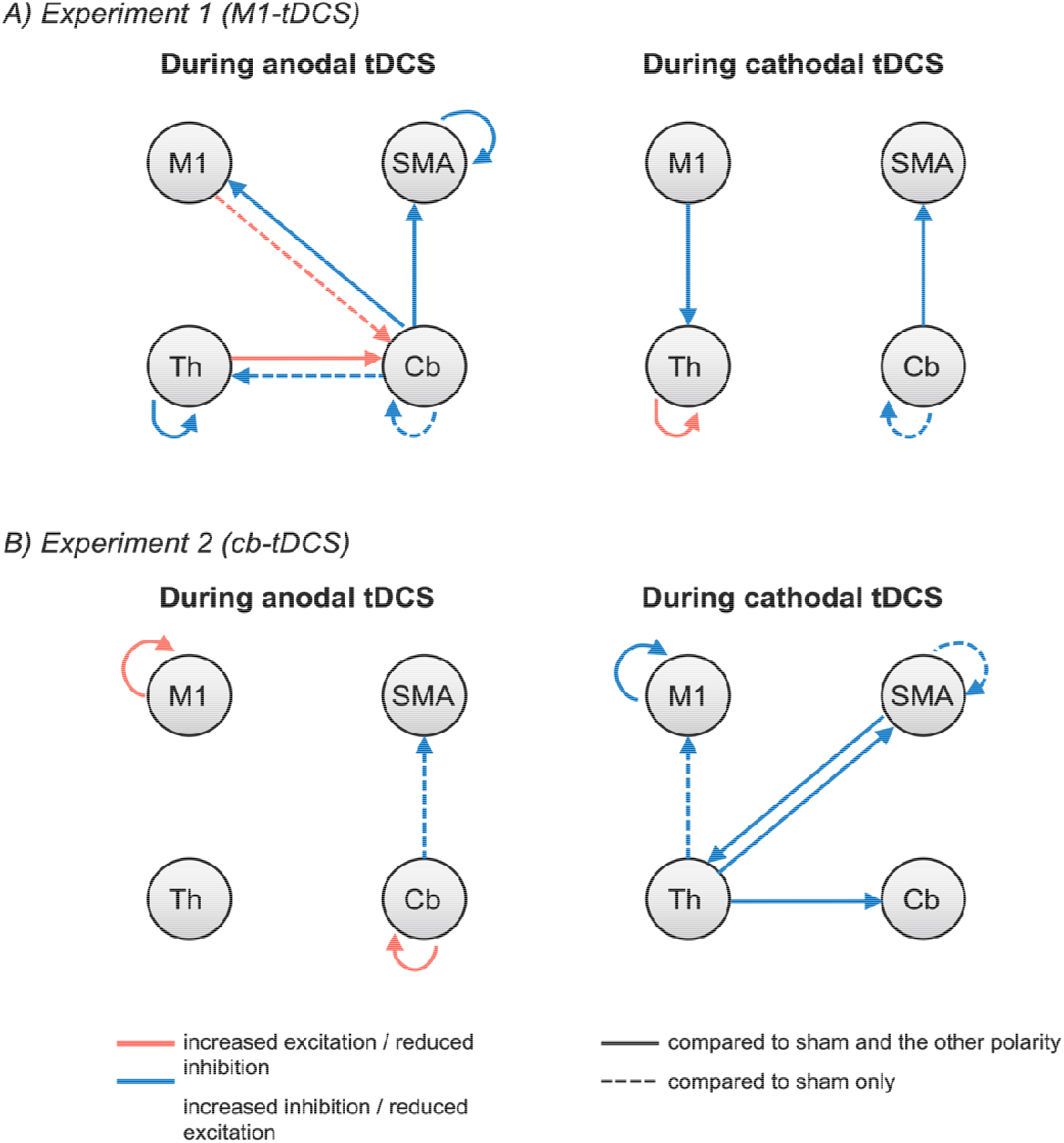
Online effects of tDCS on stationary effective connectivity. The figure shows connections that exceeded the 95% posterior probability threshold in the analysis of data during tDCS for *Experiment 1* (A) and *Experiment 2* (B). The left and right panels represent changes after anodal and cathodal stimulation respectively. Red arrows indicate connections positively related to the regressor (i.e., more excitatory / less inhibitory as a product of the comparison), while blue arrows signal connections with a negative relation to the regressor (i.e., more inhibitory/less excitatory) as a product of the comparison). Note that self-connections are always inhibitory and thus red indicates a reduction in inhibition rather than an excitatory role per-se. Solid lines represent polarity-specific effects (i.e., effects that are significant as compared to both sham stimulation and the other polarity) while dashed lines indicate significant effects as compared to sham stimulation only.

#### Experiment 2 – Effects of Cb-tDCS

The individual DCM models from this analysis explained an average of 87.18% of variance (SD = ±5.79).

During stimulation anodal and cathodal cb-tDCS respectively decreased and increased M1 self-inhibition as compared to sham and to each other. In addition, cathodal cb-tDCS increased inhibition from thalamus to cerebellum and to SMA, and from SMA to thalamus, compared to both anodal cb-tDCS and sham. When compared to sham only, cathodal tDCS also increased inhibition from thalamus to M1 as well as on the SMA self-connection.

In turn, anodal cb-tDCS reduced cerebellar self-inhibition as compared to both cathodal and sham tDCS. It also increased inhibition from cerebellum to SMA as compared to sham only. Figure 7B represents the results during cb-tDCS (see supplementary Figure S3 B for a more detailed representation).

### 3.3 Dynamic effective connectivity during tDCS

#### Experiment 1 – Effects of M1-tDCS

The individual DCM models from this analysis explained an average of 85.80% of variance (SD = ±4.16).

Focusing on the self-connections first, M1-tDCS perturbed each area differently: while anodal and cathodal tDCS appeared to slow down the rate of change in M1 and cerebellum self-connections, they increased the rate in SMA. In the thalamus self-connection, anodal tDCS was characterised by increased stability (slower rate of change), while cathodal and sham showed a similar rate with opposite sign.

Examining the extrinsic connections, we could observe that anodal tDCS slowed down the rate of change from M1 to cerebellum, while it greatly increased frequency in the connections from M1 and thalamus towards SMA, and from SMA towards thalamus and cerebellum. Cathodal tDCS instead induced higher stability (reduced rate of change) in the reciprocal connections between thalamus and cerebellum, while slightly increasing the rate of change in the connection M1 to thalamus.

Both anodal and cathodal tDCS increased the rate from cerebellum to M1 compared to sham, although maintaining different levels of connectivity strength overall.

Interestingly, both anodal and cathodal tDCS maintained weaker connectivity strength than sham in the connection from M1 to thalamus only between window 3 and 8. The two polarities also induced the same connectivity pattern from thalamus to M1 until halfway through (window 5), but started differentiating in the second half of the stimulation period (although both retained stronger connectivity than sham). These dynamics are represented in Figure 8 (top panel; see supplementary Figure S4 for the DCM results before linear combination).

**Figure 8.**
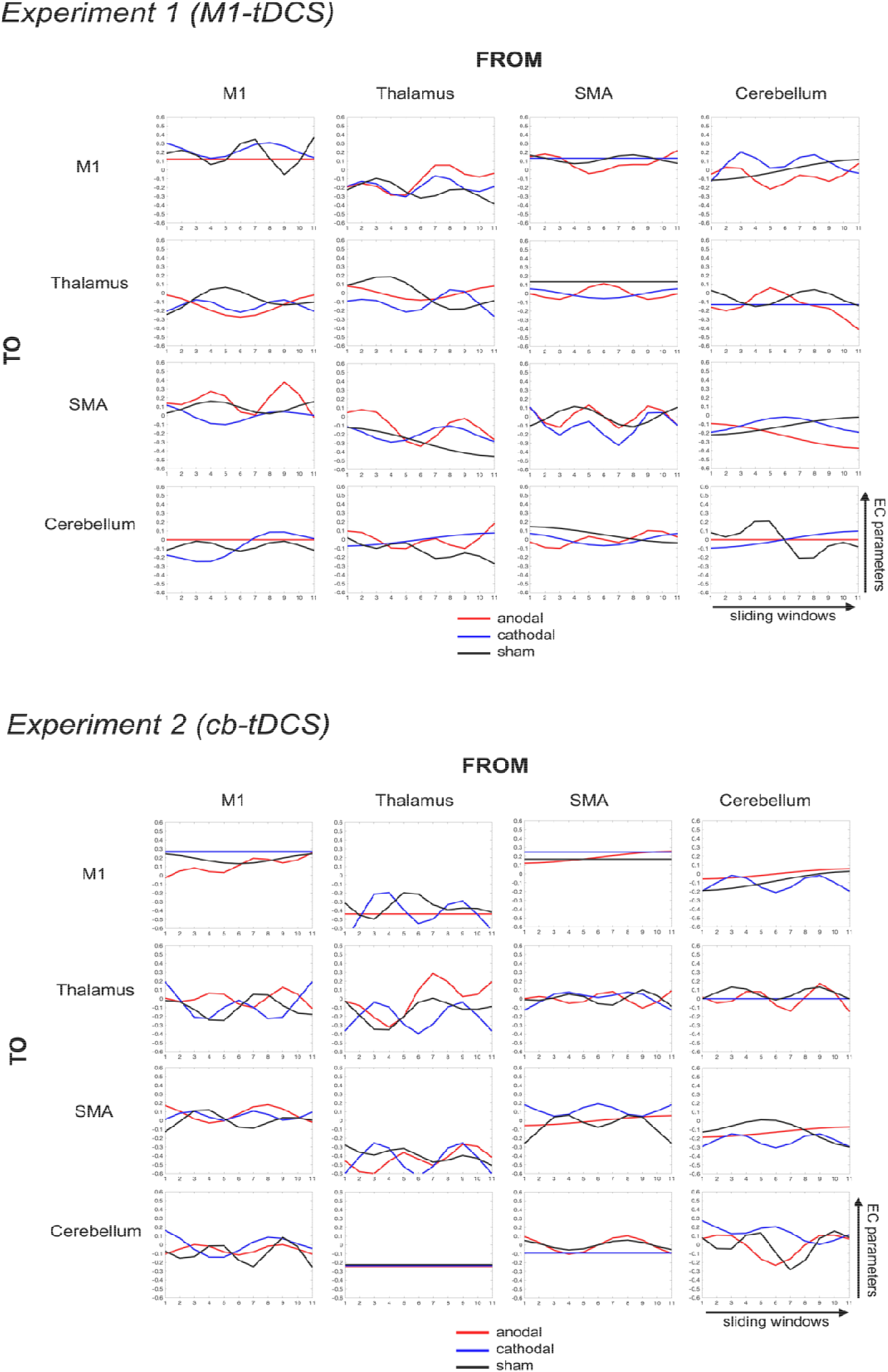
Online effects of tDCS on dynamic effective connectivity. The plots show the rate of change in effective connectivity for each connection under each tDCS polarity, calculated via linear combination of the resulting DCT basis functions weighted by the corresponding Ep parameters. Top section: Experiment 1, i.e., data during M1-tDCS. Bottom section: Experiment 2, i.e., data during cb-tDCS.

#### Experiment 2 – Effects of cb-tDCS

The individual DCM models from this analysis explained an average of 86.35% of variance (SD = ±5.20).

Similarly to M1-tDCS, cb-tDCS displayed diverse temporal characteristics in each self-connection. Interestingly, the rate of change of all polarities in the cerebellar self-connection seemed to converge to highly similar connectivity strength towards the end of stimulation, after showing ample differences halfway through stimulation (the most noticeable change being between cathodal and sham tDCS at window 5 and 6). The thalamic self-connection was instead characterised by a great differentiation between all three polarities from window 5 onwards (increased strength in anodal, decreased in cathodal tDCS). Anodal cb-tDCS increased the rate of change in the M1 self-connection (compared to the slower rates of cathodal and sham) and decreased it in SMA, where cathodal tDCS instead maintained the same rate as sham tDCS, albeit with a more inhibitory tone overall.

Overall, cb-tDCS exerted ample perturbations in some of the extrinsic connections, particularly those involving the thalamus (e.g., reciprocal connections between thalamus and M1, and from thalamus to SMA). Specifically, anodal cb-tDCS slowed the rate of change from thalamus to M1 and from cerebellum to SMA, and greatly increased it in the connection from cerebellum to thalamus. In contrast, cathodal tDCS decreased the rate from cerebellum to thalamus (as well as from SMA to cerebellum), and increased it from cerebellum to SMA and M1. These dynamics are represented in Figure 8 (bottom panel; see supplementary Figure S5 for the DCM results before linear combination).

### 3.4 Blinding

McNemar’s tests revealed no significant differences in the number of times that participants perceived active and sham tDCS as real in either experiment (Experiment 1: *X*^2^ = 11; *p* = 0.69; Experiment 2: *X*^2^ = 8; *p* = 0.38), suggesting that blinding was successful.

## 4. Discussion

In this study, we investigated the effects of tDCS over a network of four cortical and subcortical motor regions when stimulating the left M1 (Experiment 1) or right cerebellum (Experiment 2). We measured the effective connectivity of resting-state fMRI data before, during, and after tDCS (anodal, cathodal, and sham) using spectral DCM and PEB approaches. In addition, we analysed the data during tDCS using a sliding windows approach to DCM in order to display the effects of stimulation over time.

Overall, our results confirmed that the action of tDCS extends beyond the stimulated site, affecting distal connected regions in the motor network. Importantly, this included subcortical structures even when tDCS was applied over “surface” brain regions, in line with our previous study with task fMRI data (Aloi et al., 2022). Notably, we here demonstrated that both M1- and cb-tDCS are able to induce changes within the thalamo-cortico-cerebellar loops even at rest, in the absence of a motor task. Specifically, when looking at the post-stimulation period as compared to baseline, anodal M1-tDCS (Experiment 1) decreased inhibition in the thalamic self-connection, in accordance with our previous study on command following fMRI data on the same cohort (Aloi et al., 2022), and in the connection from cerebellum to thalamus. This is a key finding given that the thalamus often appears greatly inhibited in clinical conditions such as disorders of consciousness (Fernández-Espejo et al., 2010; Schiff, 2010), causing decreased cortical excitation (Fernández-Espejo et al., 2015). Being able to compensate for such thalamic hyper-inhibition could thus influence cortical activity and potentially improve motor responsiveness, for instance in the case of patients with cognitive-motor dissociation (Stafford et al., 2019). However, in contrast with our previous results during command following (Aloi et al., 2022), here we did not find other significant changes in the outputs from M1 to the rest of the network after anodal M1-tDCS. Interestingly, previous studies suggested that M1 drives the connectivity of the motor network during active movements, while having a less influential role at rest (Jiang et al., 2004), possibly explaining our results.

In turn, cathodal M1-tDCS (Experiment 1) increased inhibition from M1 to thalamus compared to both sham and anodal tDCS, in line with the canonical inhibitory role commonly attributed to this polarity. However, cathodal M1-tDCS also increased excitation across multiple connections, including that from thalamus to M1, and in agreement with Aloi et al., (2022). We have previously argued (Aloi et al., 2022) that, during movement blocks, this unexpected excitation could respond to a compensatory mechanism reacting to the inhibition received from cathodal tDCS to maintain performance at a baseline level. It is possible that a similar compensatory drive is at play here to try and re-establish the original balance in the network at rest, i.e., the baseline connectivity present at rest before perturbing the system with stimulation. Similar explanations can be found in animal studies reporting sustained homeostatic regulation of the network after stimulation, indicating a delayed re-adaptation towards baseline (pre-tDCS) levels of activation (Jackson et al., 2016; Reato et al., 2010). As the network configuration at rest is different to during task engagement, it is not surprising that the effects on specific connections do not fully match across our studies.

In Experiment 2, after applying cathodal tDCS over the cerebellum, we observed widespread effects over most connections of the network, in line with our previously reported task data (Aloi et al., 2022). However, the tone of such effects was not always in agreement. Of particular interest, here we found increased excitation from cerebellum towards cortical areas, and from thalamus to M1, in agreement with previous studies suggesting that cb-tDCS can modulate the cerebellar-brain inhibition by inhibiting the cerebellum and increasing thalamic excitation towards the cortex as a result (Galea et al., 2009). However, we also found that cathodal cb-tDCS increased inhibition from cerebellum towards the thalamus, as well as thalamic self-inhibition, both of which seem to suggest an opposite effect on CBI. While our model includes direct connections between cerebellum and cortical areas, DCM does not imply that such connections are indeed anatomically present, and instead they can be mediated by a third region (Seghier et al., 2011). In the case of CBI, there are no direct anatomical projections from cerebellum to M1 nor SMA and its influence is exerted via the thalamus (Bostan et al., 2013).

Our results may thus indicate a competing mechanism for potentiating or reducing CBI to try to maintain the balance of the network in the absence of a task.

Interestingly, anodal cb-tDCS also increased inhibition from cerebellum to thalamus, but not to the cortical areas. Such increase in inhibition was not present in our previous task results (Aloi et al. (2022), possibly indicating a stronger CBI at rest when no cortical excitation is required for motor performance (Kassavetis et al., 2011). In either case, the partial overlap in results between polarities hints at a much more complex picture than what generally observed with cortical excitability or single-neuron measures (Kwon et al., 2008). Recent simulation studies have provided a plausible explanation for the presence of similar effects across polarities, indicating a delayed increase in the connectivity within the targeted neuronal population regardless of whether tDCS induced depolarisation or hyperpolarisation during stimulation (Lu et al., 2019). Nonetheless, despite some similarities, anodal cb-tDCS mainly induced changes in connectivity towards the thalamus, whereas cathodal cb-tDCS exerted a more widespread influence across the network, including excitation from all areas towards M1.

In terms of the resting state data *during* stimulation, the connectivity patterns of ongoing M1- tDCS were different from those observed in the comparison between pre- and post-stimulation. For example, the effect of anodal M1-tDCS over the thalamic self-connection changed tone from increasing inhibition during tDCS to decreasing it after. In contrast, other connections only showed either immediate (i.e., during stimulation) effects or after-effects. For example, online anodal M1-tDCS modulated the coupling between M1 and cerebellum in both directions, and online cathodal M1-tDCS reduced thalamic inhibition, but these effects were no longer present after stimulation. This notwithstanding, some effects were consistent between the during and the post-tDCS runs. For example, ongoing anodal M1-tDCS did not seem to impact M1 or the reciprocal connections between M1 and thalamus, while ongoing cathodal M1-tDCS did increase inhibition from M1 to thalamus, both in agreement with the post-tDCS data. Similarly, the increased inhibition in the M1 self-connection after cathodal cb-tDCS already appeared during stimulation, while the effects over the connectivity from thalamus to M1 and cerebellum changed in tone between the two analyses.

Overall, in both experiments, the after-effects appeared richer (i.e., affected more connections) than the immediate effects of ongoing tDCS, possibly reflecting previously reported delayed effects of tDCS over cortical excitability, which have been linked to various types of (structural and functional) plasticity (Nitsche et al., 2004). In addition, stationary analyses average effective connectivity over the 20 minutes of ongoing stimulation, and thus more transient, incremental, and or subtle effects might be concealed. Observing how the connectivity parameters varied across time allowed us to explore the apparent discrepancies between during and after stimulation effects in more depth, as well as to understand why some predicted changes on key connections (i.e., between thalamus and M1) did not display the expected changes. For example, anodal M1-tDCS reduced the connectivity strength from M1 to thalamus halfway through stimulation (peaking at windows 5 and 6 as seen in Figure 8), but then reached a similar connectivity strength as sham towards the end of stimulation, likely explaining the absence of a stationary effect. Relatedly, an increased connection strength from thalamus to M1 was only evident in the second half of stimulation, perhaps representing a period of latency before thalamic inputs are sent towards the cortex. More broadly, cathodal M1-tDCS induced a decreased rate of change and more stable connectivity in the M1 self-connection, as compared to sham. The great instability of this connection during sham stimulation might have masked the effect of cathodal M1-tDCS in the stationary analysis, due to the averaging across the full duration of stimulation. Finally, we could observe that anodal M1-tDCS slowed down the rate of change in the thalamic self-connection and this caused and inhibitory dip that lasted for over half of the 20 minutes, with an upwards trend towards the end of the block. This might explain why we observe increased inhibition during M1-tDCS but decreased inhibition after M1-tDCS, as reported above. In contrast, cathodal M1-tDCS had a similar temporal signature to sham tDCS but with opposite phase, which could have again masked any effect in the stationary comparison. In Experiment 2 cb-tDCS influenced the rate of change in the cerebellum-to-thalamus connection, with anodal increasing and cathodal decreasing it. Relatedly, the thalamic self-connection increased its strength in the second half of anodal cerebellar stimulation, and decreased it halfway through cathodal stimulation. As above, this may explain why the related effects are observed only after stimulation but not during.

Overall, our results go against the idea of cumulative tDCS effects that get linearly stronger with longer stimulation periods, as indirectly backed-up by the studies investigating the outcomes of varying block length and interval duration (Monte-Silva et al., 2013), and support the use of dynamic approaches to gain a richer understanding of nuanced effects on spatiotemporal connectivity patterns. Previous research has indeed suggested a nonlinearity between tDCS parameters and after-effects, and called for more nuanced experimental approaches (Batsikadze et al., 2013; Jamil et al., 2020; Monte-Silva et al., 2013). Understanding these dynamic modulations could lead to optimised tDCS protocols that can tailor the montage and duration to maximise the desired effects on the target connections (Filmer et al., 2020).

There are a number of further considerations to be acknowledged. First, in our first Experiment, we observed broad effects on SMA and its connections (especially after anodal M1-tDCS), which could indicate current dispersion from the nearby target electrode. Previous studies on current modelling indeed showed that conventional tDCS is characterised by low spatial resolution, whereby the current can spread on the scalp beyond the target area due to skull and tissue dispersion (Datta et al., 2009). While this constitutes a wider issue for the neurostimulation community, it calls for caution when interpreting our M1-tDCS data in terms of causality between direct effects over M1 and the broader changes in the network. Furthermore, the choice of the window size and overlap in dynamic connectivity analyses is non-trivial and remains an arbitrary decision (Preti et al., 2017). In our study we selected the shape, length, overlap and number of windows to match the study from Park and colleagues as closely as possible (Park et al., 2018). However, due to difference in the TR between our respective datasets (2 and 2.7 seconds in Experiments 1 and 2, and 0.72s in Park et al., 2018), we could not exactly replicate the window parameters, and our windows included fewer time-points (100 instead of 200), precluding a direct comparison between studies. Nevertheless, the specific effects of window parameters on effective connectivity are yet to be established. Similarly, we employed a DCT with 6 functions following (Park et al., 2018) and as is particularly indicated for modelling resting-state dynamics. However, the optimal set of temporal basis functions broadly, but also specifically to modelling tDCS changes, also requires further investigation.

## 5. Conclusions

Overall, our results show broad spatiotemporal modulations of the motor network both during and after stimulation for both M1- and cb-tDCS. These effects extended beyond the stimulated site, inducing widespread changes on distant (including subcortical) structures. Some of these changes were immediate and took place during stimulation, while others became evident only after the stimulation ended. Our analysis of dynamic connectivity shed light into the underpinning mechanisms for these apparent discrepancies and allowed us to observe that the rate of change of each connection was influenced by tDCS in a unique manner.

Our results call into question the traditional view of anodal and cathodal tDCS as clearly excitatory and inhibitory, respectively, as well as the idea of a cumulative effect of tDCS over time. Indeed, they reveal a fuller and more intricate picture at the network level. Our more nuanced approach may provide useful information for optimising the use of tDCS for clinical and cognitive purposes with all the montages.

## CRediT authorship contribution statement

Sara Calzolari: Formal analysis, Methodology, Writing – original draft, Writing – review & editing. Roya Jalali: Investigation. Davinia Fernández-Espejo: Visualization, Funding acquisition, Data curation, Formal analysis, Methodology, Supervision, Writing – review & editing.

## Data availability statement

Processed data is available from the authors upon reasonable request. Please contact with any questions or requests.

## Acknowledgments

This work was supported by generous funding from the Medical Research Council (MR/P02596X/1; DF-E). SC was supported by a studentship from the Biotechnology and Biological Sciences Research Council (BBSRC) Midlands Integrative Biosciences Training Partnership (MIBTP).

