## Supplementary Material for "Characterising stationary and dynamic effective connectivity changes in the motor network during and after tDCS"

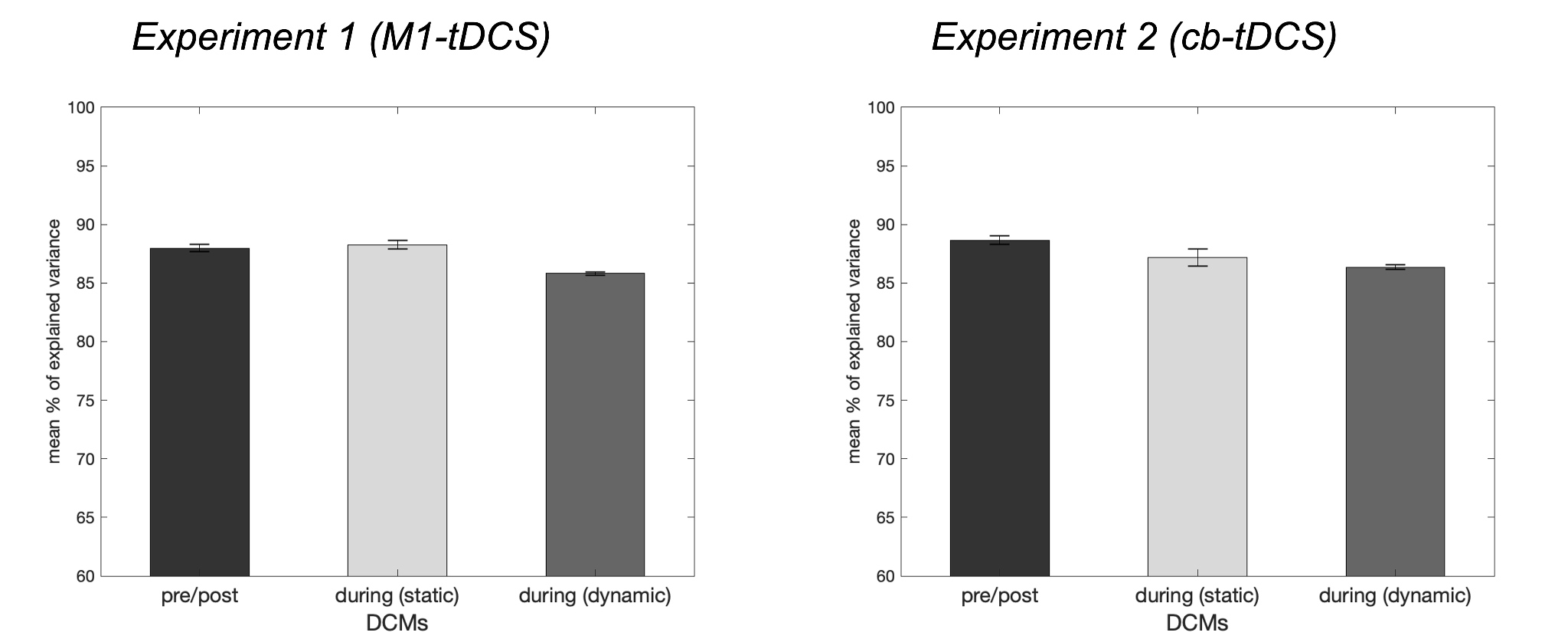

**Figure S1.** Mean percentage of variance explained by individual DCM models for each of the 3 DCM analyses (left: Experiment 1; right: Experiment 2). The error bars show the standard error of the mean.

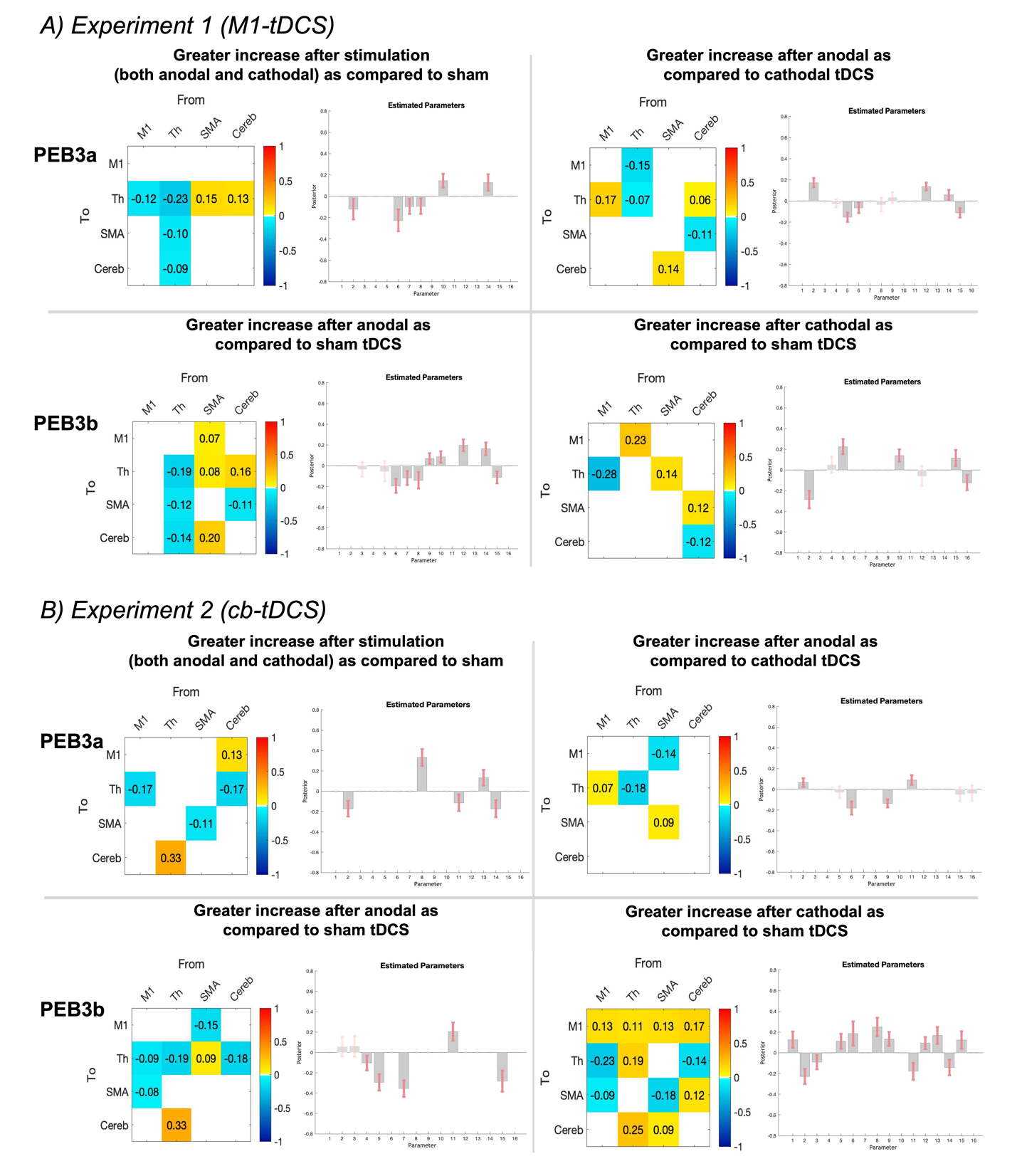
**Figure S2.** Connections that exceeded the 95% posterior probability threshold in the analysis of data after tDCS compared to baseline (interactions between time and polarity only). A) Experiment 1, i.e., results after M1-tDCS. B) Experiment 2, i.e., results after cb-tDCS. In both A and B, the top half of the figure shows results from the first PEB-of-PEBs design (PEB3a), the bottom half results from the second PEB-of-PEBs (PEB3b). In the matrices, warm colours and positive values (estimated parameters) indicate the connection is positively related to the regressor (i.e., the connection becomes more excitatory as a product of the comparison), while cold colours and negative values signal a negative relation to the regressor (i.e., the connection becomes more inhibitory as a product of the comparison). This applies to extrinsic connectivity, while self-connections are log-scaled and therefore must always be interpreted as inhibitory (cold colours signal reduced inhibition in relation to the regressor, warm colours signal increased inhibition). The bar charts show the effective connectivity parameters in grey, with the red error bars showing their relative posterior probability. Darker bars represent the parameters that surpassed the 95% posterior probability threshold.

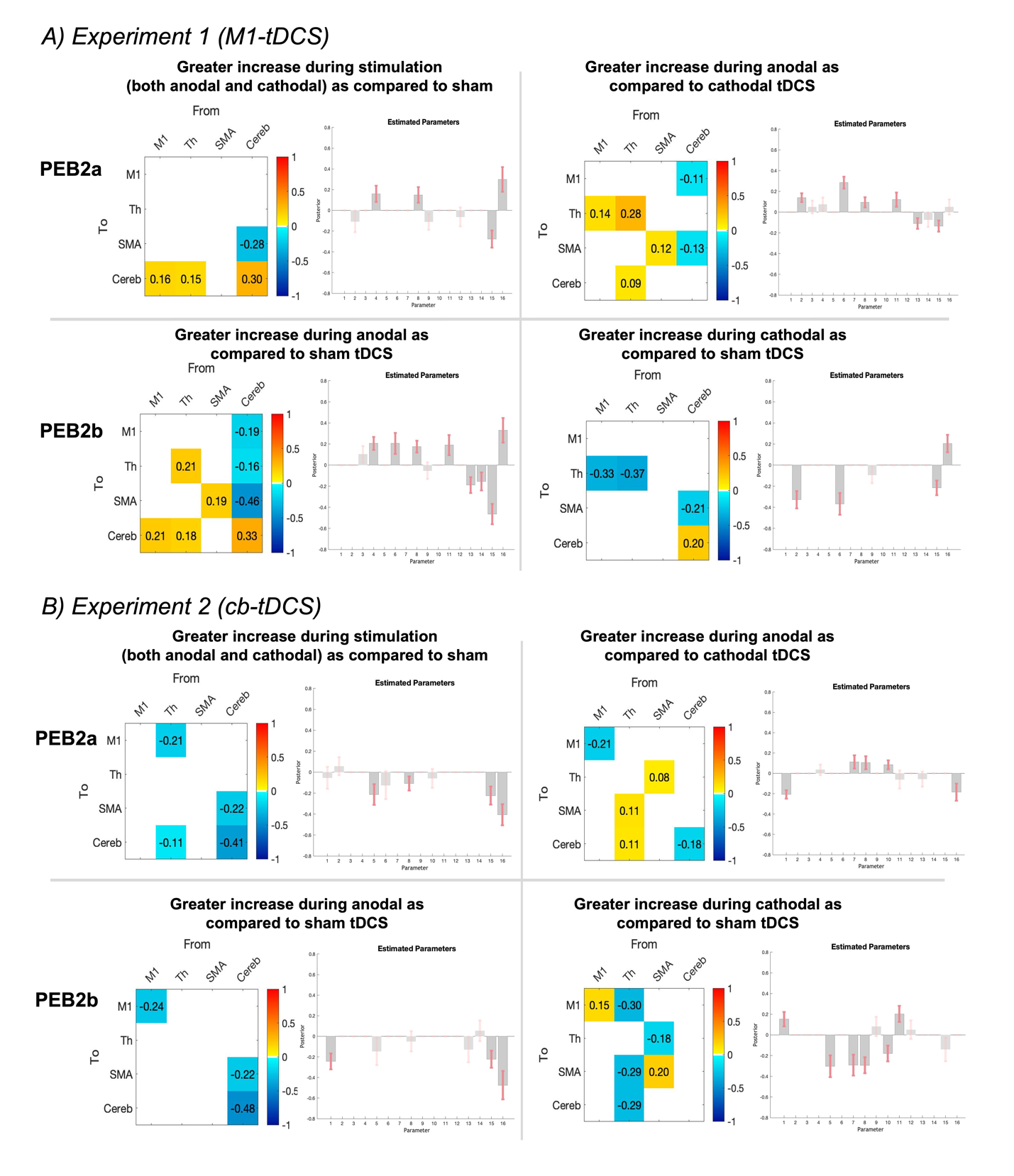

**Figure S3.** Connections that exceeded the 95% posterior probability threshold in the stationary analysis of data during tDCS. A) Experiment 1, i.e., stationary results during M1-tDCS. B) Experiment 2, i.e., stationary results during cb-tDCS. In both A and B, the top section shows results from the first PEB-of-PEBs design (*PEB2a,* testing for the effects of real tDCS compared to sham and for differences between anodal and cathodal tDCS). The bottom section shows the results from the second PEB-of-PEBs (*PEB2b*: anodal and cathodal tDCS compared to sham). In the matrices, warm colours and positive values (estimated parameters) indicate the connection is positively related to the regressor (i.e., the connection becomes more excitatory as a product of the comparison between polarities), and vice versa for cold colours and negative parameter values. This applies to extrinsic connectivity, while self-connections are log-scaled and must always be interpreted as inhibitory. The bar charts show the effective connectivity parameters in grey, with the red error bars showing their relative posterior probability. Darker bars represent the parameters that surpassed the 95% posterior probability threshold.

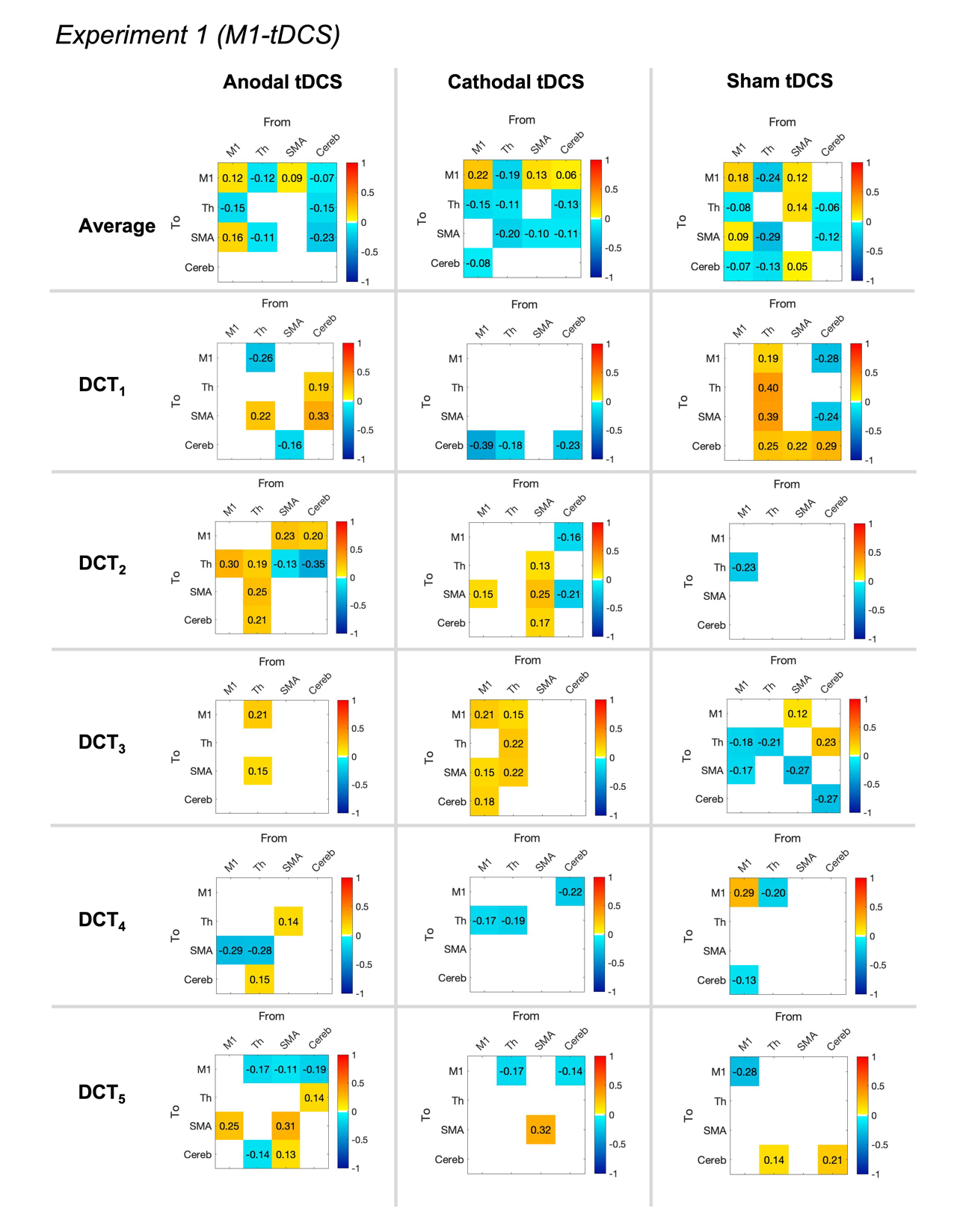

**Figure S4.** Connections that exceeded the 95% posterior probability threshold in the dynamic analysis of data during M1-tDCS (Experiment 1). The figure shows results from the second PEB-of-PEBs (*PEB2,* testing for the group effects of each tDCS polarity). In the matrices, warm colours and positive values (estimated parameters) indicate the connection is positively related to the regressor (i.e., the connection fits that specific discrete cosine function), and vice versa for cold colours and negative parameter values (i.e., the connection fits the discrete cosine function but with opposite sign).

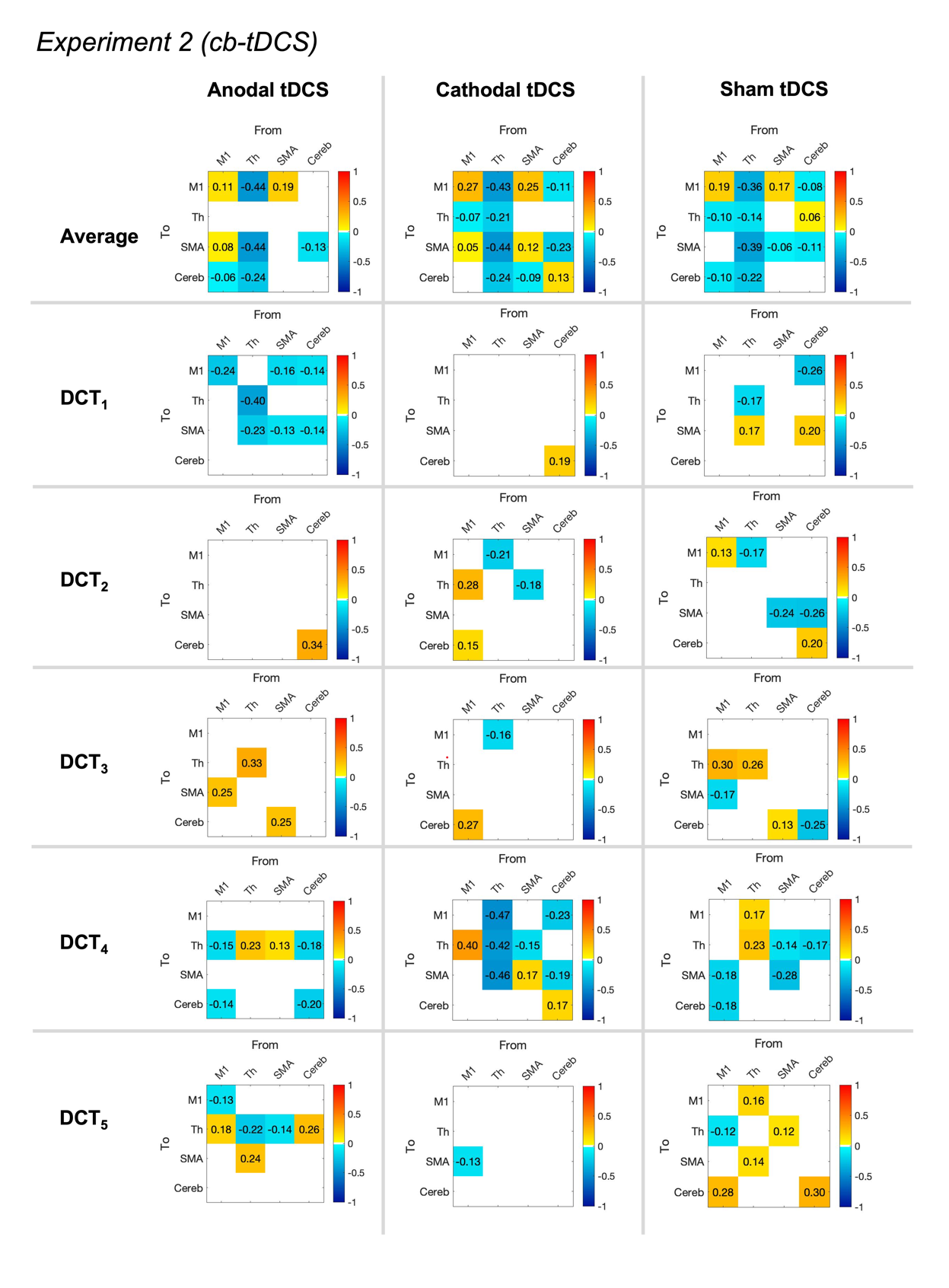

**Figure S5.** Connections that exceeded the 95% posterior probability threshold in the dynamic analysis of data during cb-tDCS (Experiment 2). The figure shows results from the second PEB-of-PEBs (*PEB2,* testing for the group effects of each tDCS polarity). In the matrices, warm colours and positive values (estimated parameters) indicate the connection is positively related to the regressor (i.e., the connection fits that specific discrete cosine function), and vice versa for cold colours and negative parameter values (i.e., the connection fits the discrete cosine function but with opposite sign).
